# Comprehensive analysis of mRNA poly(A) tail reveals complex and conserved regulation

**DOI:** 10.1101/2021.08.29.458068

**Authors:** Yusheng Liu, Hu Nie, Yiwei Zhang, Falong Lu, Jiaqiang Wang

## Abstract

Non-templated poly(A) tails are added to the 3′-end of most mRNAs, which have important roles in post-transcriptional regulation^1–3^. Recent studies have revealed that poly(A) tails are not composed purely of A residues, but also contain U, C and G residues internally and at their 3′-ends^4–6^, revealing new levels of complexity. However, no method is able to analyze these internal and terminal non-A residues simultaneously. Here, we developed a new method called unbiased capture of 3-terminal followed by full-length RNA sequencing (UCTF-seq) which captures RNA 3′-ends by direct 3′ adaptor ligation and rRNA removal by CRISPR/Cas9. This method allows simultaneous evaluation of the poly(A) tail length and 5′-end, internal, and 3′-end non-A residues together with the full-length cDNA for a transcript. Applying this method, we achieved the first complete transcriptome-wide 3′ tail map of mRNA within the nuclear and cytoplasmic compartments of mammalian cells, uncovering differences in poly(A) tail length and non-A residues between these two mRNA populations. A survey of diverse eukaryotic species revealed the conservation of a subset of poly(A) tails containing consecutive U residues in the internal positions, whereas those with consecutive C or G residues were of much lower abundance. Together, we established the first method to be able to comprehensively analyze poly(A) tail 5′-end, internal and 3′-end non-A residues in addition to the length simultaneously, and reveal the first complete mRNA 3′ tail map, providing rich insights into the regulatory roles of poly(A) tails.

## Introduction

Most mature messenger RNAs (mRNAs) and long noncoding RNAs (lncRNAs) undergo non-templated 3′ polyadenylation to form poly(A) tails. mRNA poly(A) tails have been known as essential structural components of mRNAs with important roles in mRNA stability and translation^2, 7–13^. The regulatory roles of mRNA poly(A) tails have long been underestimated partly due to the technical difficulty in analyzing poly(A) tails because of their homopolymeric nature which is problematic for second-generation sequencing platforms^4, 14^.

To overcome the difficulty in genome-wide analysis of RNA poly(A) tails, several methods were recently developed to accurately measure their length, including TAIL-seq, PAL-seq, FLAM-seq and PAIso-seq^4–6, 14^. In addition, Non-A residues in poly(A) tails (non-A residues hereafter) were found to be wide-spread^4–6^, and they are associated with a new layer of post-transcriptional regulation^4–6^. For example, U residues can be found at the 3′-end of very short poly(A) tails, and they promote the degradation of the corresponding transcripts^15^. On the contrary, G residues at the 3′-end of poly(A) tails can inhibit the enzymatic activity of deadenylase complexes, and thereby stabilize the corresponding transcript^16^. Besides accurate length quantification, PAIso-seq can accurately measure each nucleotide in the poly(A) tails. This approach revealed wide- spread non-A residues within the body of poly(A) tails, consistent with findings obtained using PacBio-based FLAM-seq^5^.

However, neither PAIso-seq nor FLAM-seq methods can measure non-A residues at the 3′-end of poly(A) tails^5, 6^, while TAIL-seq cannot measure non-A residues within the body of poly(A) tails^4^. Therefore, although non-A residues in poly(A) tail are of great interest in terms of dynamic post-transcriptional regulation, there is no currently available method for transcriptome-wide measurement of 3′-end and internal non-A residues of poly(A) tails simultaneously.

In this study, we developed a new method named unbiased capture of 3-terminal followed by full-length RNA sequencing (UCTF-seq) to achieve the first comprehensive poly(A) tail analysis method without any bias toward base composition of RNA 3′-ends. Using this method, we reveal three different patterns of U, C and G residues within poly(A) tails. Moreover, we analyzed the nuclear and cytoplasmic fractions of mRNAs and found that they differed in mRNA poly(A) tail length and non-A residue composition. In addition, we identified consecutive U residues but not G or C residues in the internal body of poly(A) tails in all the samples of representative eukaryotic species analyzed. Together, we presented a most comprehensive poly(A) tail analyzing method for the complete 3′ tail map, and reveal that non-A residues are of specific patterns and distinct subcellular features, suggesting a global new layer of non-A residue mediated post-transcriptional regulation in addition to the length of poly(A) tails.

## Results

### UCTF-seq for accurate poly(A) tail analysis

Current poly(A) tail analysis methods are not comprehensive. PAIso-seq method and FLAM-seq method both have bias toward underestimate the terminal non-A residues due to the requirement of terminal A nucleotide for efficient base-pairing mediated strand annealing^5, 6^. To address this problem, we developed a new method called UCTF- seq in which a 3′-end adaptor is directly ligated to the poly(A) tail (Fig. 1a). This strategy, which is not affected by 3′-end base composition, consequently preserves all poly(A) tail information. In addition, the ligated adaptor sequence is well defined, and can be accurately positioned for downstream data analysis. After ligation the RNA is then reverse transcribed with a primer complementary to the 3′ adaptor. The 5′-end of the cDNA is tagged through template switching using an oligo with a unique molecular identifier (isoTSO-UMI)^17^. The unbiased capture of all RNA 3′-ends leads to one problem that rRNA, the abundance of which is about 20 times more than that of the mRNA within a cell, is also ligated with the 3′-end adaptor. To efficiently remove rRNA without sacrificing the sensitivity of the method, a post-amplification rRNA depletion approach is used. Amplified cDNA from rRNA is destroyed by the combination of Cas9 protein with a pool of sgRNAs targeting rRNA cDNA sequence (Fig. 1a, Supplementary Table 1). After rRNA removal, the cDNA is amplified by a second round of PCR and used to generate a circular SMRTbell library for PacBio sequencing in HiFi mode.

**Fig. 1.**
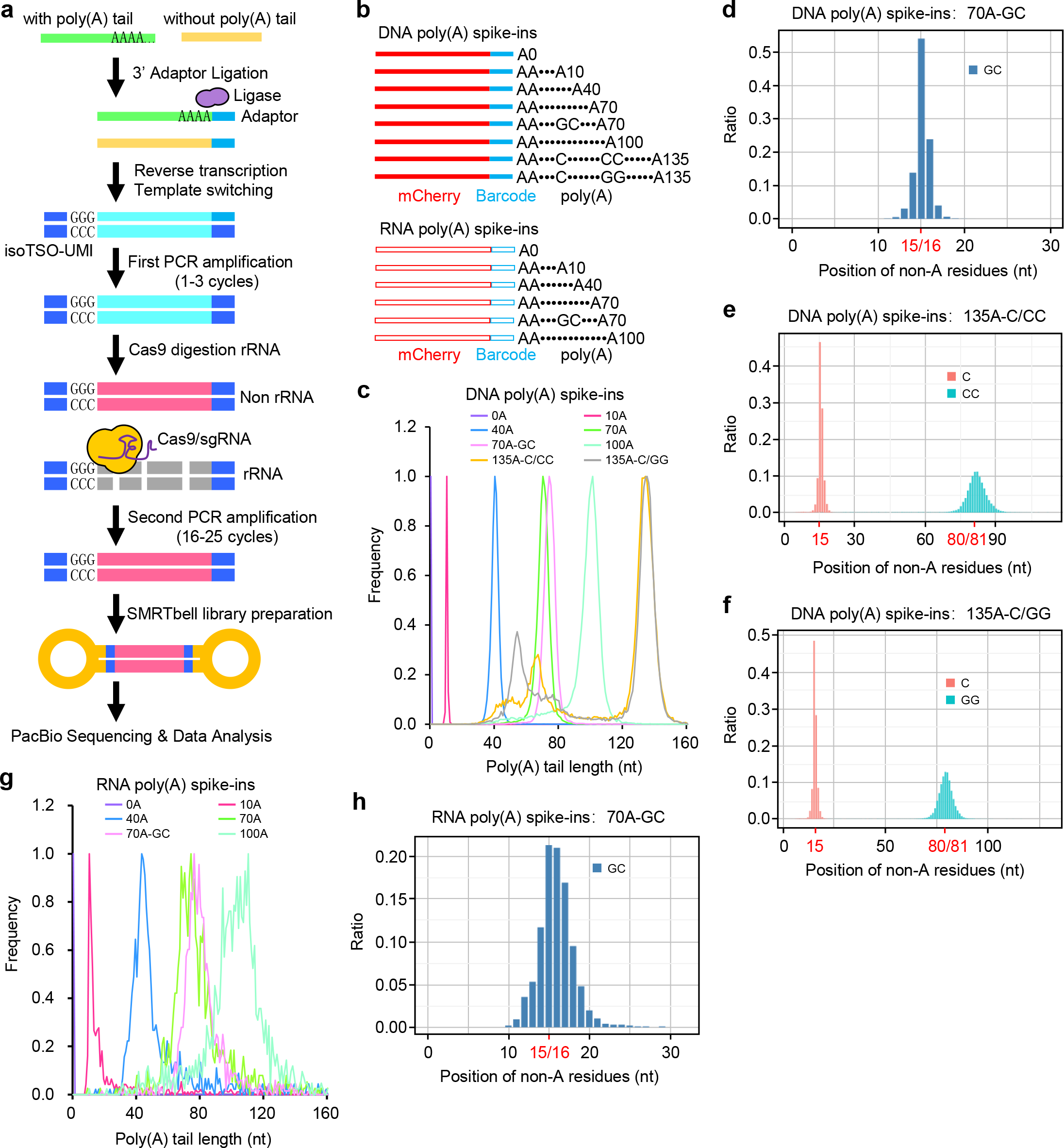
The principle and validation of UCTF-seq. **a,** Flowchart for UCTF-seq method design. The main steps of the method include 3′ adaptor ligation, reverse transcription, template switching, full length cDNA amplification, rRNA removal by CRISPR/Cas9, circular adaptor ligation and PacBio sequencing. **b,** The structure of the DNA poly(A) spike-ins (top) and RNA poly(A) spike-ins (bottom). **c,** Histogram of poly(A) length for each DNA spike-in with 1 nt bin size. The detected median length is close to the expected one (0, 10, 40, 70, 74, 99, 131, and 131 nt, respectively). The Y-axis is normalized by the maximum value of each histogram. **d - f,** Proportion of measured non-A residues at the indicated positions in the 70A-GC (**d**), 135A-C/CC (**e**), and the 135A-C/GG (**f**) DNA spike-ins. The number in red represents the expected position of the non-A residues in the standard. **g,** Histogram of poly(A) length for each RNA spike-in with 1 nt bin size. The detected median length is close to the expected one (0, 13, 46, 75, 79, and 103 nt, respectively). The Y-axis is normalized by the maximum value of each histogram. **h,** Proportion of measured non-A residues at the indicated positions in the 70A-GC RNA spike-ins. The number in red represents the expected position of the non-A residues in the standard.

To validate the performance of UCTF-seq, we sequenced a pool of barcoded synthetic cDNA and mRNA spike-ins with different poly(A) tail lengths (Supplementary Table 2). These spike-ins contain the *mCherry* gene sequence, poly(A) tails of different lengths each with a specific barcode, and a PCR handle after the poly(A) tail (Fig. 1b, Extended Data Fig. 1a and Supplementary Table 2). The mRNA spike-ins were prepared by *in vitro* transcription of the linearized DNA spike-ins. We sequenced these spike-ins into UCTF-seq runs with three replicates each and a similar number of reads was obtained for each of the replicates (Extended Data Fig. 1b, c). We observed sharp peaks at or very close to the expected length for the DNA spike-ins (Fig. 1c), similar to previous observations using both PAIso-seq and FLAM-seq analysis^5, 6^. For the 135 nt tails, secondary peaks of smaller size were observed (Fig. 1c), which might represent recombination of long homopolymers which is known to happen in *E. coli*^18, 19^. As non-A residues were observed within the body of poly(A) tails, we also included non-A residues within the DNA spike-ins to evaluate the performance of UCTF-seq in sequencing them. The non-A residues were identified at the expected positions (Fig. 1d-f), suggesting this approach can accurately characterize non-A residues within poly(A) tails.

For the IVT mRNA spike-ins, we observed peaks close to the expected size although the peaks were not as sharp as the DNA spike-ins (Fig. 1g). The reduced resolution of peaks seen here for IVT mRNA spike-ins compared to the DNA spike-ins is likely due to the inaccuracy of IVT in synthesis of the desired length of poly(A) tail from DNA templates with defined poly(A) tail lengths. Indeed, LC-MS analysis of IVT poly(A) tails revealed a relatively wide distribution of poly(A) tail length rather than a single length^20^. In addition, the non-A residues within the mRNA poly(A) spike-ins were detected around the expected sites (Fig. 1h). Together, these results confirm that UCTF- seq can accurately measure both the length and non-A residues of the poly(A) tails.

### Systematic comparison of poly(A) tail sequencing methods

In addition to the new UCTF-seq method, there are several poly(A) analyzing methods available, including PAL-seq^14^, PAL-seq2^21^, TAIL-seq^4^, mTAIL-seq^22^, PAT-seq^23^ and TED-seq^24^ on the Illumina platform, FLAM-seq^5^ and PAIso-seq^6^ on the PacBio platform, and Nanopore Direct RNA-seq on the ONT platform^25^. Comparing the features of these methods, we can see that UCTF-seq is the only method that can analyze the complete sequence features of RNA poly(A) tails, including length, terminal non-A residues, and internal non-A residues simultaneously (Fig. 2a). In addition, UCTF-seq together with PAIso-seq, FLAM-seq, and Nanopore Direct RNA-seq can be used to analyze the poly(A) tail as well as the full-length RNA isoform (Fig. 2a).

**Fig. 2.**
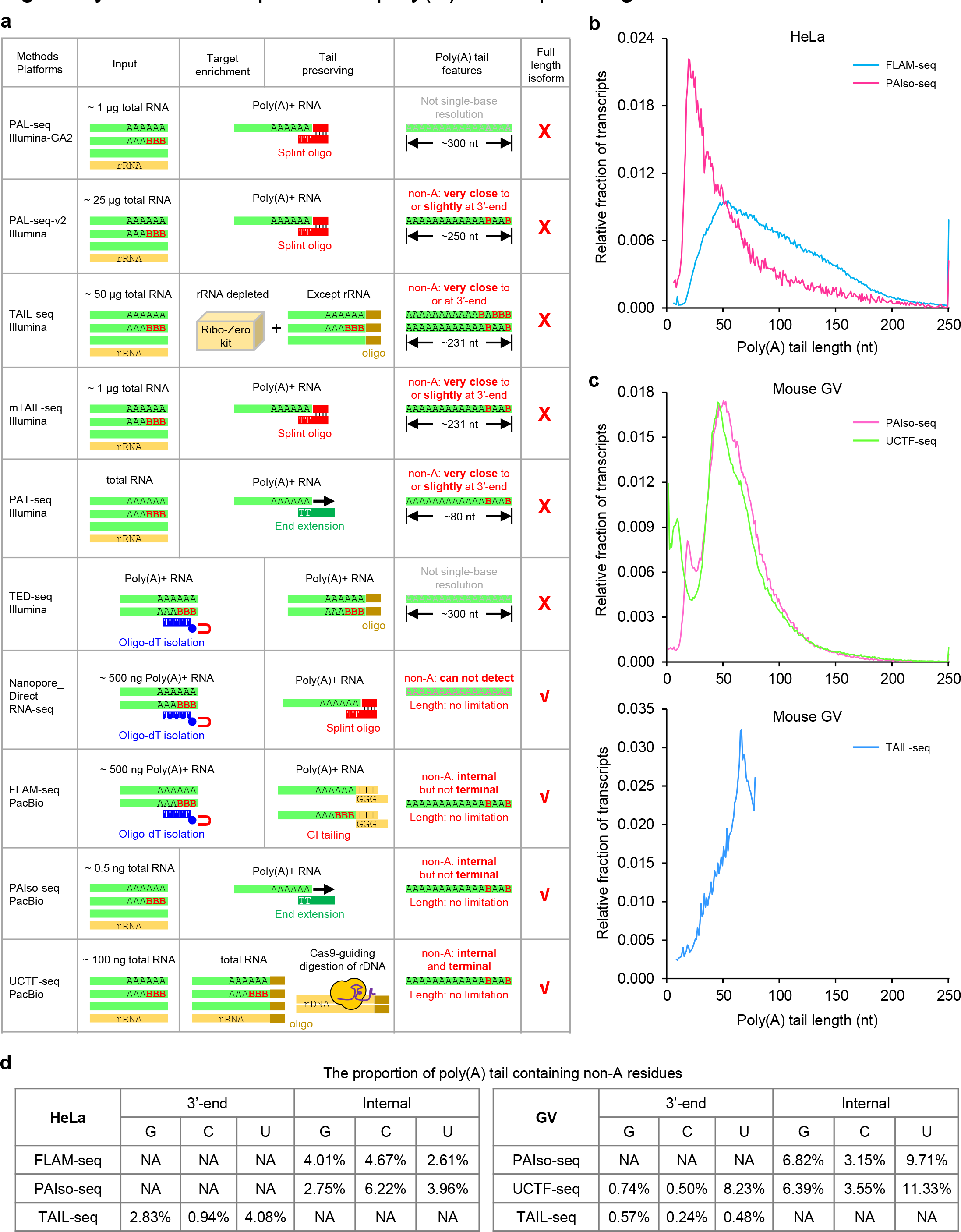
Systematic comparison of poly(A) tail sequencing methods. **a,** Table summarizing the key features of the methods, including sequencing platform, maternal input requirement, mRNA enrichment method, tail preserving method, feature of the output data, and with full-length isoform information or not. **b,** PAIso-seq comparison with FLAM-seq showing the advantage of no requirement of poly(A)+ RNA enrichment by oligo-dT beads, which introduces bias towards long poly(A) tails. Data shown here are acquired from HeLa cells: PAIso-seq (this study), FLAM-seq^5^. Histograms (bin size = 1 nt) are normalized to cover the same area. Transcripts with poly(A) tail of at least 1 nt are included in the analysis. Transcripts with poly(A) tail lengths greater than 250 nt were included in the 250 nt bin. **c,** UCTF-seq outperforms PAIso-seq in capturing the fraction of transcripts of very short poly(A) tails (top). TAIL-seq has a major drawback in the upper-limit of the tail length detection. All the data shown here are acquired from mouse GV oocyte samples: PAIso-seq^6^, TAIL-seq^26^ and UCTF-seq (this study). Histograms (bin size = 1 nt) are normalized to cover the same area. Transcripts with poly(A) tail of at least 1 nt are included in the analysis. Transcripts with poly(A) tail length greater than 250 nt are included in the 250 nt bin. **d,** Proportion of transcripts containing non-A residues in HeLa cells detected by FLAM- seq^5^, TAIL-seq^4^, and PAIso-seq^6^, and in GV oocytes detected by PAIso-seq^6^, UCTF- seq and TAIL-seq^26^. UCTF-seq is the only method to be able to measure non-A residues at 5′-end, Internal and 3′-end parts simultaneously.

UCTF-seq and PAIso-seq do not require a poly(A)+ RNA selection step, which avoids the bias toward long poly(A) tails. The bias introduced by poly(A)+ RNA enrichment is evident in the FLAM-seq data^5^ when compared to PAIso-seq data for the same HeLa cells (Fig. 2b). In addition, we performed UCTF-seq analysis of mouse GV oocytes, and compared the data to two published datasets generated using PAIso-seq^6^ or TAIL-seq^26^. A very similar pattern was found among the poly(A) tails longer than 25 nt resulted from the PAIso-seq and UCTF-seq data (Fig. 2c), but UCTF-seq captured a much greater number of tails < 20 nt compared to PAIso-seq (Fig. 2c). Notably, both UCTF-seq and PAIso-seq methods outperform TAIL-seq in analyzing the length of poly(A) tails in GV oocytes (Fig. 2c). The biggest drawback of this TAIL-seq dataset is the upper-limit of 79 nt in measuring poly(A) tails which makes it unable to measure the length a substantial proportion of poly(A) tails correctly.

In addition to the poly(A) tail length, we looked into the non-A residues detected in the above datasets. Using FLAM-seq and PAIso-seq datasets, similar proportion of poly(A) tails were detected with internal non-A residues, but the non-A residues at the 3′-end of poly(A) tails could not be quantified (Fig. 2d). In contrast, TAIL-seq data could be used to quantify the 3′-end non-A residues of poly(A) tails, but not to quantify the internal non-A residues (Fig. 2d). As expected, the new UCTF-seq method can be used to quantify both 3′-end and internal non-A residues of poly(A) tails (Fig. 2d). Taken together, the above data demonstrate that UCTF-seq is useful for discovery and accurate measurement of both length and non-A residues of mRNA poly(A) tails.

### Non-A residues from 5′-end to 3′-end of poly(A) tails

With evident advantages in analyzing mRNA poly(A) tails, we used UCTF-seq to investigate the pattern of non-A residues distribution in the mRNA poly(A) tails of 3T3 cells (3T3-T: 3T3 total RNA) and mouse embryonic stem cells (ES cells) (ES-T: ES total RNA). The results revealed good correlations between replicates on gene expression level and poly(A) tail length (Extended Data Fig. 2a, b). In addition, the UCTF-seq data showed a good coverage of full-length transcripts, further demonstrating the good quality of the data (Extended Data Fig. 2c, d). Distribution of non-A residues for different lengths of poly(A) tails ranging from 1 nt to 180 nt revealed the presence of U, C, and G residues at the 5′- and 3′-ends, and internal positions of poly(A) tails (Fig. 3a, b, d). One common distribution pattern observed was enrichment of the U, C, and G residues at the very 5′-end of poly(A) tails, but U and G residues showed stronger 3′-end enrichment than C residues (Fig. 3a, b).

**Fig. 3.**
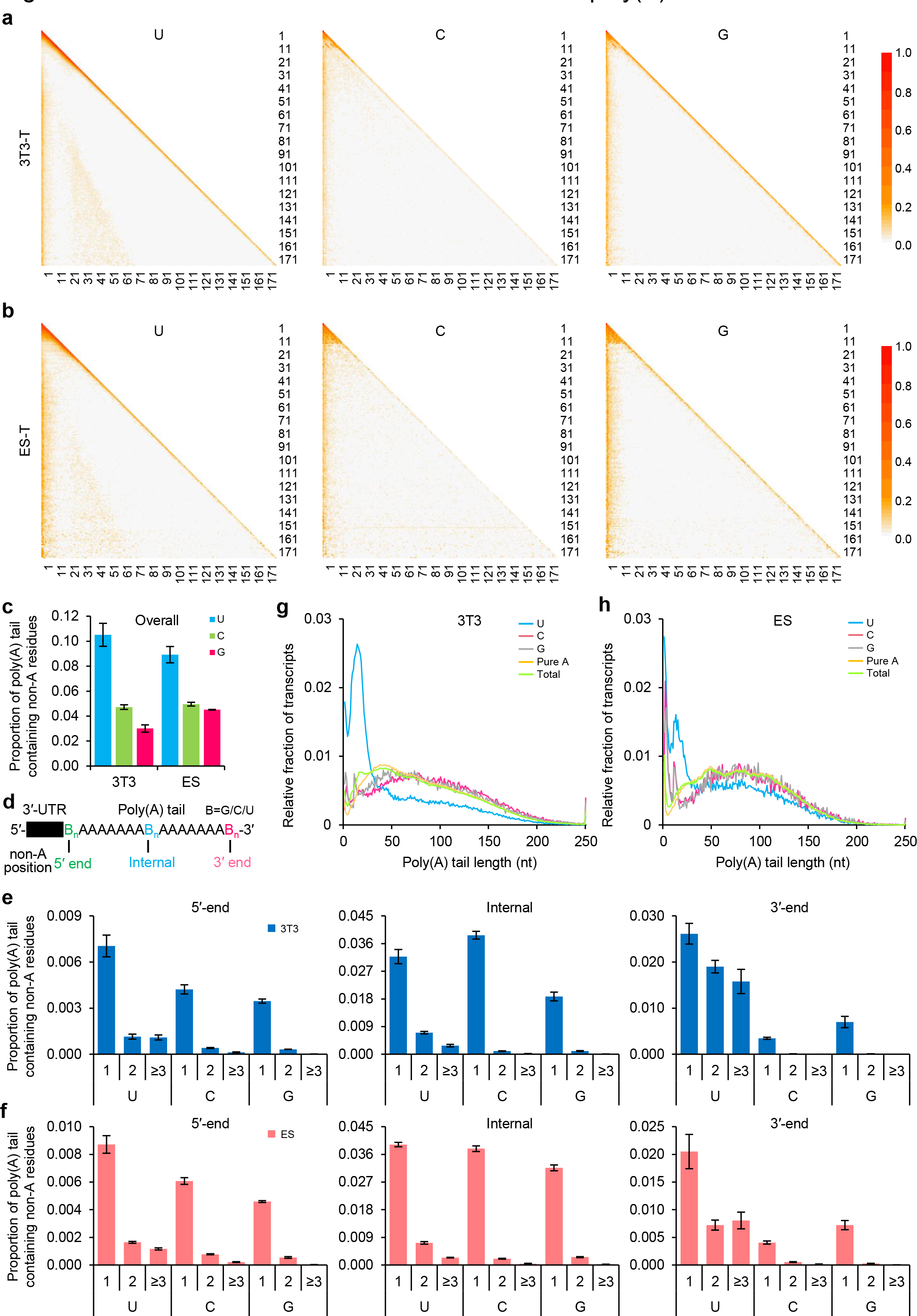
The overall distribution of non-A residues in the poly(A) tails. **a, b,** Distribution patterns of U, C or G residues within poly(A) tails in 3T3-T (3T3 total RNA) (**a**) and ES-T (ES total RNA) (**b**) samples. Poly(A) tails with indicated non-A residues of a given length are collapsed to one line. The relative abundance of non-A residues at each position is calculated and visualized by a color scale. Poly(A) tails with length between 1-180 nt are included and ranked in the heatmap from 1-180 (top to bottom). **c,** Overall proportion of transcripts containing U, C and G residues in 3T3 and ES samples. Transcripts with poly(A) tails of at least 1 nt are included in the analysis. The ratio of reads containing U, C or G for is calculated by the number of reads containing U, C or G residues divided by the total number of reads with poly(A) tails of at least 1 nt. Error bars indicate the standard error of the mean (SEM) from two replicates. **d,** Diagram of the positions (5′-end, Internal and 3′-end) of non-A residues. **e, f,** Proportion of transcripts containing U, C and G residues at 5′-end (left), internal (middle) and 3′-end (right) positions in 3T3 (**e**) or ES (**f**) samples. The non-A residues are further divided according to the length of the longest consecutive U, C or G (1, 2, and ≥ 3). Transcripts with poly(A) tails of at least 1 nt are included in the analysis. The ratio of reads containing the indicated number of U, C or G residues is calculated by the number of reads containing U, C or G residues divided by the total number of reads with poly(A) tails of at least 1 nt. Error bars indicate the SEM from two replicates. **g, h,** Histogram of lengths of total transcripts, transcripts with U, C or G residues, and transcripts composed of pure A sequence in 3T3 (**g**) or ES (**h**) cells. Histograms (bin size = 1 nt) are normalized to cover the same area. Transcripts with poly(A) tails of at least 1 nt are included in the analysis. Transcripts with poly(A) tail lengths greater than 250 nt are included in the 250 nt bin.

Overall, in both 3T3 and ES cells, the frequency of transcripts harboring U, G, or C residues was 10, 4, and 4 percent, respectively (Fig. 3c). At the individual gene level, the poly(A) tail length and percentage of poly(A) tails containing non-A residues was also similar between the two types of cells (Extended Data Fig. 3, Supplementary Table 3). Frequencies of poly(A) tails with non-A residues at their very 5′-end, 3′-end, or internally (non-A residues that were separated by at least one A residue from both ends) were determined (Fig. 3e, f). As previously reported, the non-A residues were sometimes present as singletons or as consecutive oligos in the poly(A) tails^4, 6^. Therefore, we investigated the distribution of single, double and oligo (three or more consecutive) non-A residues at the three different positions in poly(A) tails (Fig. 3e, f). C and G were found to exist largely as single residues, whereas U was often present as a doublet or longer stretch (Fig. 3e, f). No position effects were seen. Examination of the length of the poly(A) tails bearing non-A residues revealed that the tails with U residues are relatively short, while tails with C and G residues are slightly depleted in the short fraction (Fig. 3g, h). Taken together, these data reveal that U, C and G residues are present at the 5′ and 3′ ends of poly(A) tails, as well as internally, which suggests potential regulatory roles.

### mRNA poly(A) tails contain different levels of non-A residues in the nucleus and cytoplasm

We then investigated potential differences in poly(A) tails characteristics between nuclear and cytoplasmic mRNA, a question that has not been previously explored. Cytoplasmic and nuclear mRNA was separately isolated from 3T3 and ES cells (3T3- N: 3T3 nuclear RNA, 3T3-C: 3T3 cytoplasmic RNA, ES-N: ES nuclear RNA, ES-C: ES cytoplasmic RNA), following a previously published method^27, 28^. Then we performed UCTF-seq with these samples (Extended Data Fig. 2). By analyzing the distribution of known nuclear or cytoplasmic enriched RNA in our UCTF-seq data, we confirmed that our nuclear-cytoplasmic RNA separation was successful (Extended Data Fig. 4a, b). For example, *Neat1* and *Malat1,* well known nuclear speckle localized lncRNAs^28, 29^, are enriched in the nuclear RNA fraction, whereas coding mRNAs are enriched in the cytoplasmic fraction (Extended Data Fig. 4a, b). Additionally, the mitochondrial transcripts are very little in the nuclear fraction, further confirming the successful separation of nuclear and cytoplasmic RNA (Extended Data Fig. 4c).

**Fig. 4.**
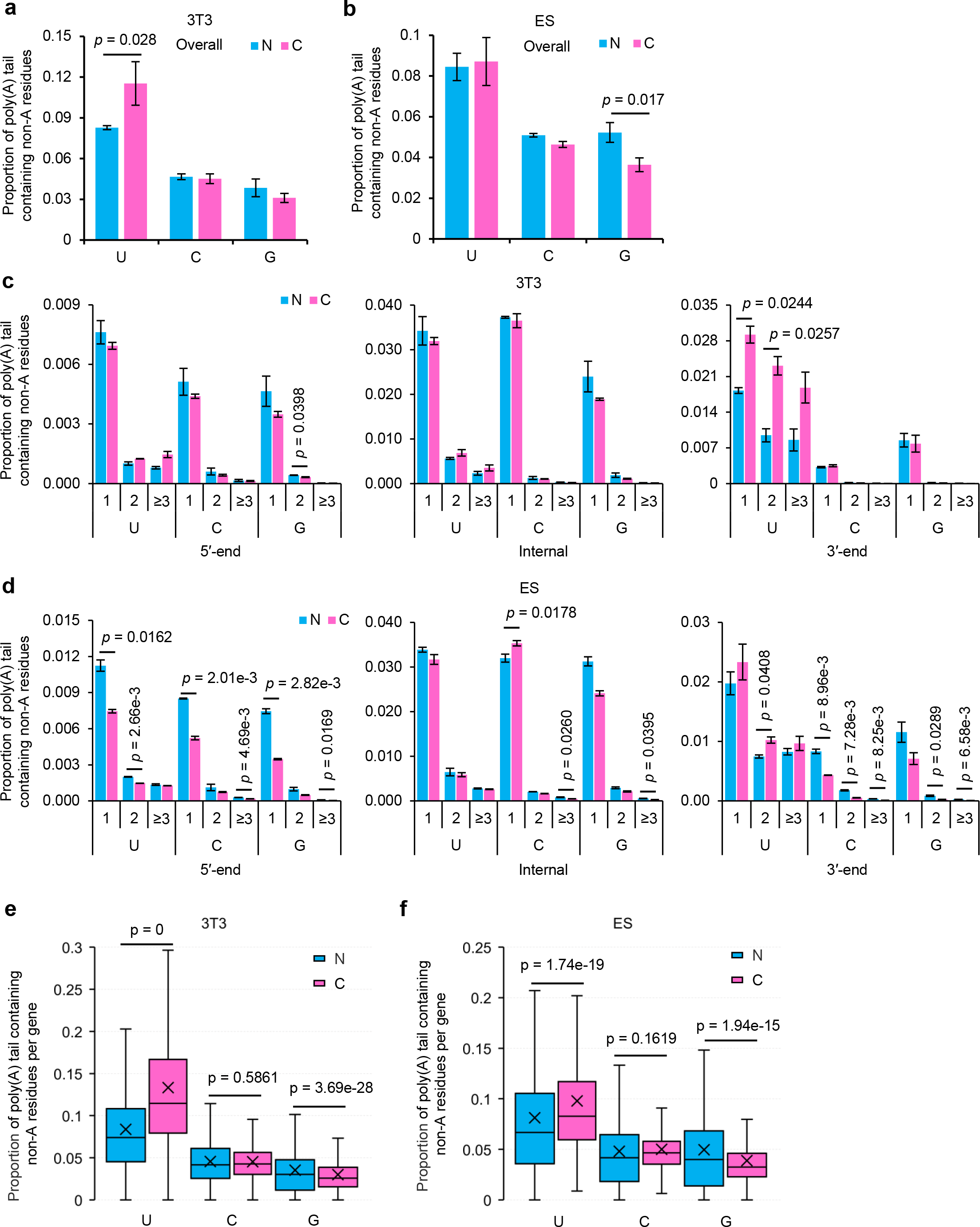
Comparison of non-A residues between the nuclear and the cytoplasmic fractions. **a, b,** Overall proportion of transcripts containing U, C or G residues within poly(A) tails in the 3T3 (**a**) or ES (**b**) nuclear (N) and the cytoplasmic (C) fractions. Error bars indicate the SEM from two replicates. **c, d,** Proportion of transcripts containing U, C or G residues at 5′-end (left), internal (middle) and 3′-end (right) positions in the 3T3 (**c**) or ES (**d**) nuclear (N) and the cytoplasmic (C) fractions. The non-A residues were further divided according the length of the longest consecutive U, C or G (1, 2, and ≥ 3). Error bars indicate the SEM from two replicates. **e, f,** Box plot of the proportion of reads containing U, C or G residues for each gene in the 3T3 (**e**) or ES (**f**) nuclear (N) and the cytoplasmic (C) fractions. Genes (n = 4,756 for 3T3, n = 1,165 for ES) with at least 20 poly(A) tail containing reads (tail length ≥ 1) for both N and C fractions are included in the analysis. The “×” indicates the mean value, black horizontal bars show the median value, and the top and bottom of the box represent the values of the 25^th^ and 75^th^ percentiles, respectively. Transcripts with poly(A) tails of at least 1 nt are included in the analysis. The ratio of reads containing U, C or G residues is calculated by dividing the number of reads containing U, C or G residues by the total number of reads with poly(A) tails of at least 1 nt. The differences between the N and C fractions are tested by Student′s *t* test. *P* values are shown for comparisons that are statistically significant.

Examination of the positions of non-A residues in the poly(A) tails of nuclear and cytoplasmic mRNAs revealed a similar overall distribution pattern (Extended Data Fig. 5). However, a significantly higher level of overall U residues was observed in the cytoplasmic fraction of 3T3 cells (Fig. 4a) and significantly lower level of overall G residues was found in the cytoplasmic fraction of ES cells (Fig. 4b). Looking into different positions, overall, cytoplasmic mRNAs contained more U residues at the 3′- ends, and fewer U residues at the 5′-ends of poly (A) tails compared with nuclear mRNAs (Fig. 4c, d). In contrast, the levels of G and C residues present at the 5′-end, internal, and 3′-ends in poly(A) tails are generally lower in the cytoplasm compared to those in the nucleus (Fig. 4c, d). These trends were generally reflected in poly(A) tails of transcripts of individual genes, with the proportion of U residues increasing, the proportion of C residues being minimally affected, and the proportion of G residues decreasing in the cytoplasmic vs nuclear fractions (Fig. 4e, f, Supplementary Table 3). The differing distributions of U, C and G residues within mRNA poly(A) tails in the nuclear and cytoplasmic mRNA fractions, suggests a differential mechanism of synthesis of these residues and potentially distinct functions of the U, C and G residues in the nucleus and cytoplasm.

**Fig. 5.**
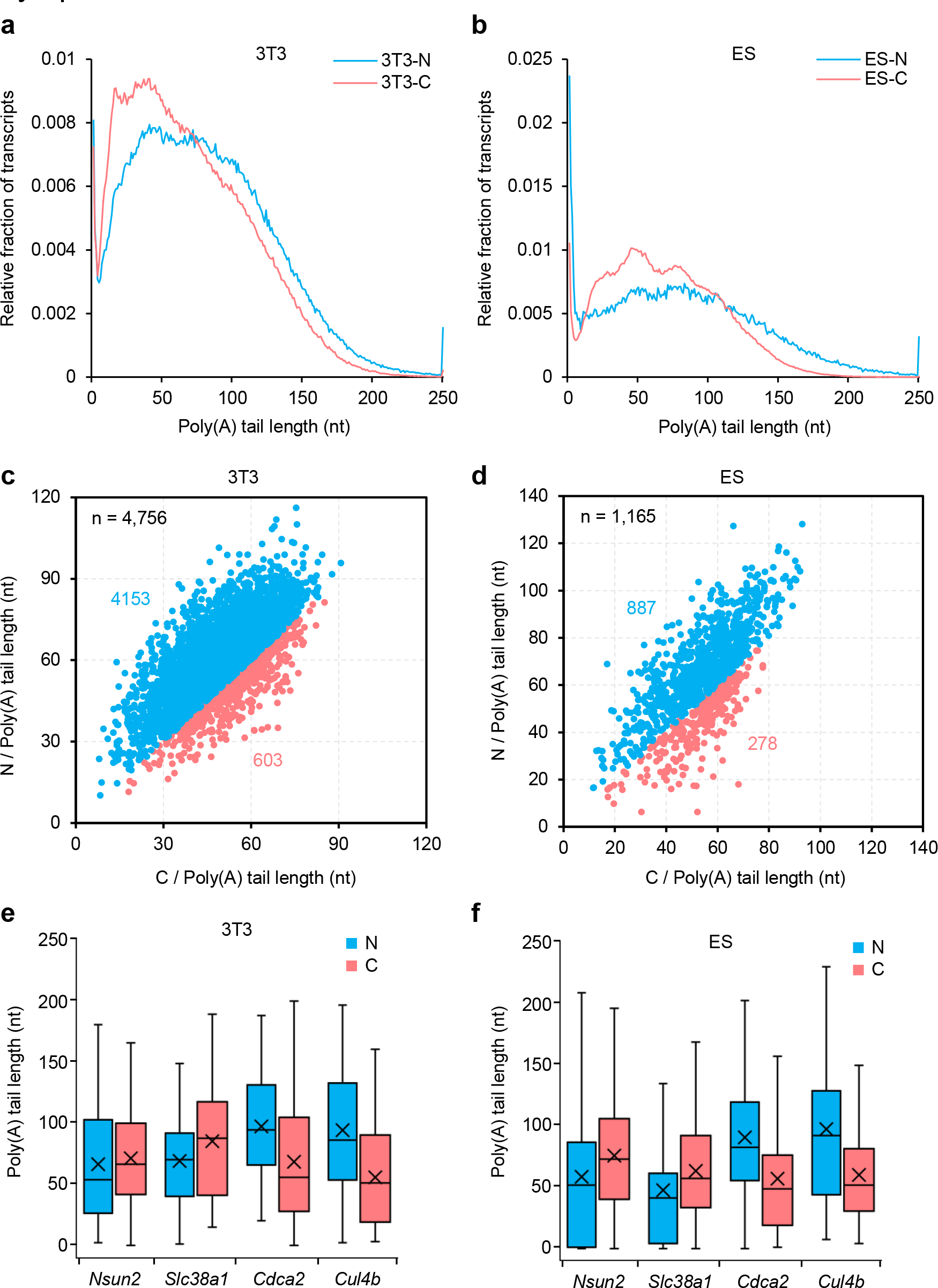
Poly(A) tail length is generally longer in the nuclear fraction compared with the cytoplasmic fraction. **a, b,** Global distribution of poly(A) tail lengths of all transcripts in the 3T3 (**a**) or ES (**b**) nuclear (N) and the cytoplasmic (C) fractions. Histograms (bin size = 1 nt) are normalized to cover the same area. Transcripts with poly(A) tails of at least 1 nt are included in the analysis. Transcripts with poly(A) tail lengths greater than 250 nt were included in the 250 nt bin. **c, d,** Scatter plot of poly(A) tail lengths for each gene in the 3T3 (**c**) or ES (**d**) nuclear (N) and the cytoplasmic (C) fractions. Each dot represents one gene. Dots in blue are genes with longer poly(A) tails in the nuclear fraction, while dots in red are genes with longer poly(A) tails in the cytoplasmic fraction. The poly(A) tail length for each gene is the geometric mean length of all the transcripts with poly(A) tails of at least 1 nt for the given gene. Genes with at least 20 reads in each of the replicates are included in the analysis. The number of genes included in the analyses is indicated at the top left of the graphs. **e, f,** Box plot for the poly(A) tail length of *Nsun2*, *Slc38a1*, *Cdca2* and *Cul4b* in the 3T3 (**e**) or ES (**f**) nuclear (N) and the cytoplasmic (C) fractions. Transcripts with poly(A) tail of at least 1 nt for the given gene are included in the analysis. The “×” indicates mean value, the black horizontal bars show the median value, and the top and bottom of the box represent the value of 25^th^ and 75^th^ percentile, respectively.

### mRNA poly(A) tails are of different length in the nucleus and cytoplasm

Cleavage and polyadenylation are known to be essential for export from the nucleus to the cytoplasm^30^. Blocking mRNA nuclear export has also been linked to hyperadenylation of some specific transcripts^31, 32^. However, it is not clear whether nuclear mRNA simply have longer poly(A) tails or if the blockade of mRNA export leads to a hyperadenylation response. Our data provide an answer to this question. The nuclear transcripts of 3T3 and ES cells were found to have obviously longer poly(A) tails than cytoplasmic ones (Fig. 5a, b). This phenomenon was also seen in individual genes; for the majority of genes the poly(A) tails of nuclear transcripts was longer than those of cytoplasmic transcripts, and a small number of genes showed the opposite phenomenon (Fig. 5c, d, Supplementary Table 4). For example, *Cdca2* and *Cul4b* showed shorter poly(A) tails, while *Nsun2* and *Slc38a1* show longer poly(A) tails in the cytoplasm compared to that in the nucleus in both cell types (Fig. 5e, f). Therefore, these data reveal that for most genes, poly(A) tails are longer in the nucleus than those in the cytoplasm, suggesting important regulation and function of poly(A) tails in these compartments.

### Conserved consecutive U residues within the body of mRNA poly(A) tails in eukaryotes

Consecutive U residues are present in the internal body of poly(A) tails in all the 3T3 cell, ES cell and GV oocyte^6^ samples, whereas the consecutive C or G residues are less frequent. Therefore, the poly(A) tails containing internal consecutive U residues may represent a class of poly(A) tails with distinct biochemical properties and functions. To investigate this, we first asked whether the consecutive U residues were present in mRNA from other tissues and other species. We analyzed RNA from different types of mouse organs, including brain, epididymis, heart, intestine, kidney, liver, lung, muscle, ovary, oviduct, spleen, testis, uterus, white adipose tissue (WAT), and brown adipose tissue (BAT) (Fig. 6a), as well as total RNA from several representative organisms including *Arabidopsis thaliana*, *C. elegans*, fish, fly, human HeLa cells, pig, rat, rice and yeast (Fig. 6b). Interestingly, we can see that there are various levels of non-A residues within the poly(A) tails in all these samples (Fig. 6a, b). In particular, similar to what we observed in 3T3 and ES cell lines, consecutive U residues were often found (Fig. 6c, d), while C and G residues were identified mainly as single nucleotide residues (Fig. 6c, d).

**Fig. 6.**
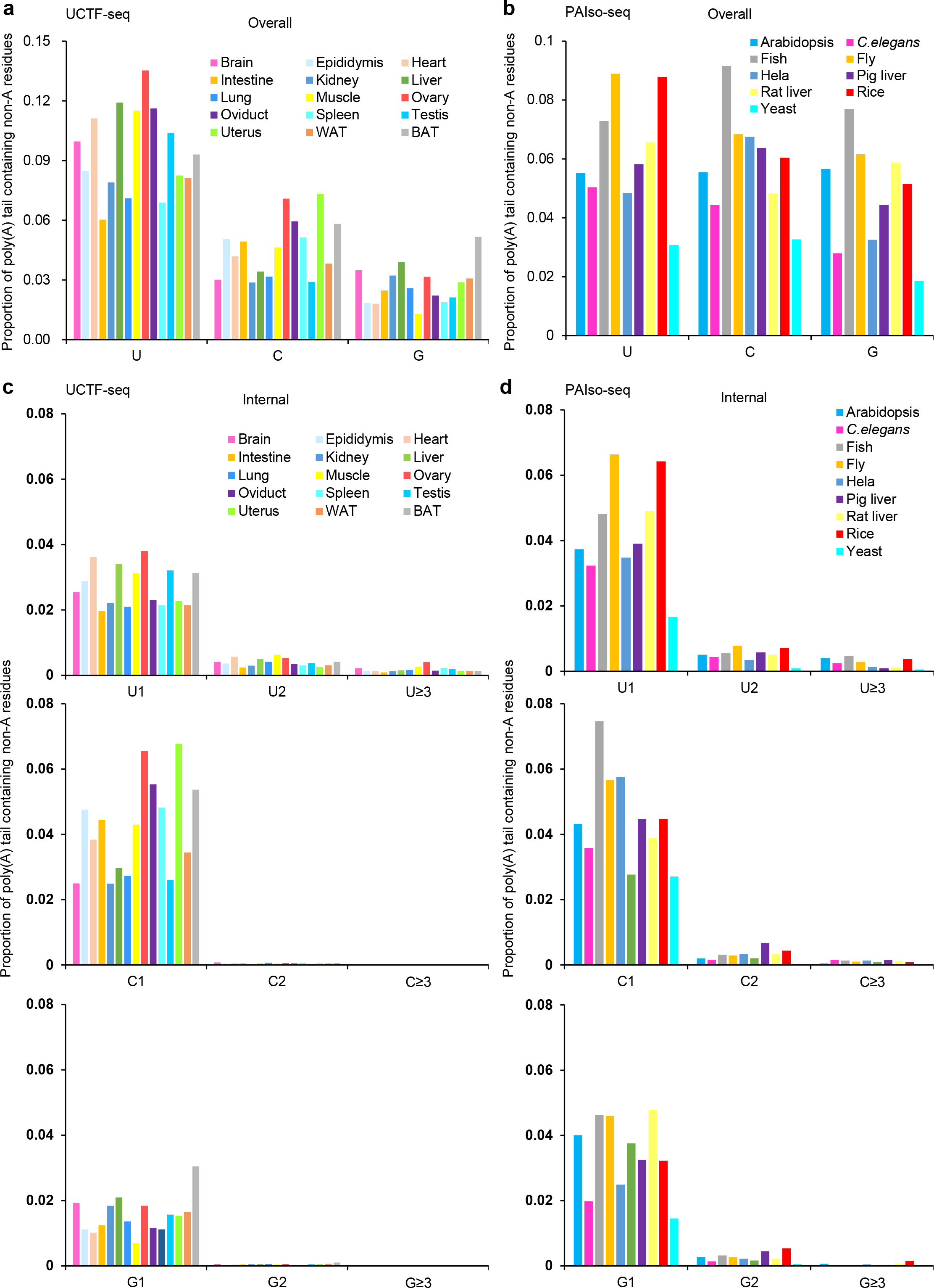
Internal non-A residues in poly(A) tails are conserved across eukaryotes. **a,** Overall proportion of transcripts containing U, C or G residues within poly(A) tails in different mouse tissue. WAT: white adipose tissue, BAT: brown adipose tissue. The data here are generated using UCTF-seq. **b,** Overall proportion of transcripts containing U, C or G residues within poly(A) tails in different model organisms. The data here are generated using PAIso-seq. **c, d,** Proportion of transcripts containing U (top), C (middle) or G (bottom) residues in internal body of poly(A) tails in different mouse tissue as revealed by UCTF-seq (**c**) or in different model organisms as revealed by PAIso-seq (**d**). The non-A residues was further divided according to the length of the longest consecutive U, C or G (1, 2, and ≥ 3). WAT: white adipose tissue, BAT: brown adipose tissue. The data here are generated using UCTF-seq. Transcripts with poly(A) tails of at least 1 nt are included in the analysis. The ratio of reads containing U, C or G for is calculated by the number of reads containing U, C or G residues divided by the total number of reads with poly(A) tails of at least 1 nt.

Taken together, these data reveal that transcripts with poly(A) tails having internal consecutive U residues are highly conserved among different eukaryotic species and may therefore represent a distinct subclass, with potentially important biological roles.

## Discussion

mRNA poly(A) tails have long been known to be one of the most important structural components of mRNA in eukaryotes. Traditionally, mRNA poly(A) tails were considered to be composed of a pure stretch of A residues, the presence of which is essential for the stability and initiation of translation^33^. However, it has become clear that the regulatory roles of poly(A) tails have been highly underestimated. For a long time, the length of the poly(A) tail was the only factor thought to be involved in dynamic regulation of mRNA fate and function (Fig. 7a). The discovery of non-A residues at the 3′-end of mRNA poly(A) tails introduced other regulatory possibilities^7^. Indeed, 3′ U residues have been implicated in accelerating degradation of short tailed mRNAs^15^, while the 3′ G residues have been suggested to stabilize poly(A) tails by inhibiting the enzymatic activity of deadenylase complexes^16^. The recently developed PAIso-seq and FLAM-seq methods together revealed the existence of internal non-A residues within mRNA poly(A) tails (Fig. 7a). These discoveries raised the question whether the presence of non-A residues in poly(A) tails is widespread and whether these residues have biological functions. However, none of the existing methods were able to simultaneously measure non-A residues in both internal and 3′-end poly(A) tails. In this study, we developed a new method, UCTF-seq, to achieve the complete 3′ tail map of the transcriptome. Very interestingly, we found that non-A residues can occur at the 5′- end, internal, and 3′-end parts of poly(A) tails (Fig. 7a) across diverse species and tissues. The specific distribution patterns of U, C or G residues within poly(A) tails suggest that they can carry out specific highly regulated biological functions, which await further investigation.

**Fig. 7.**
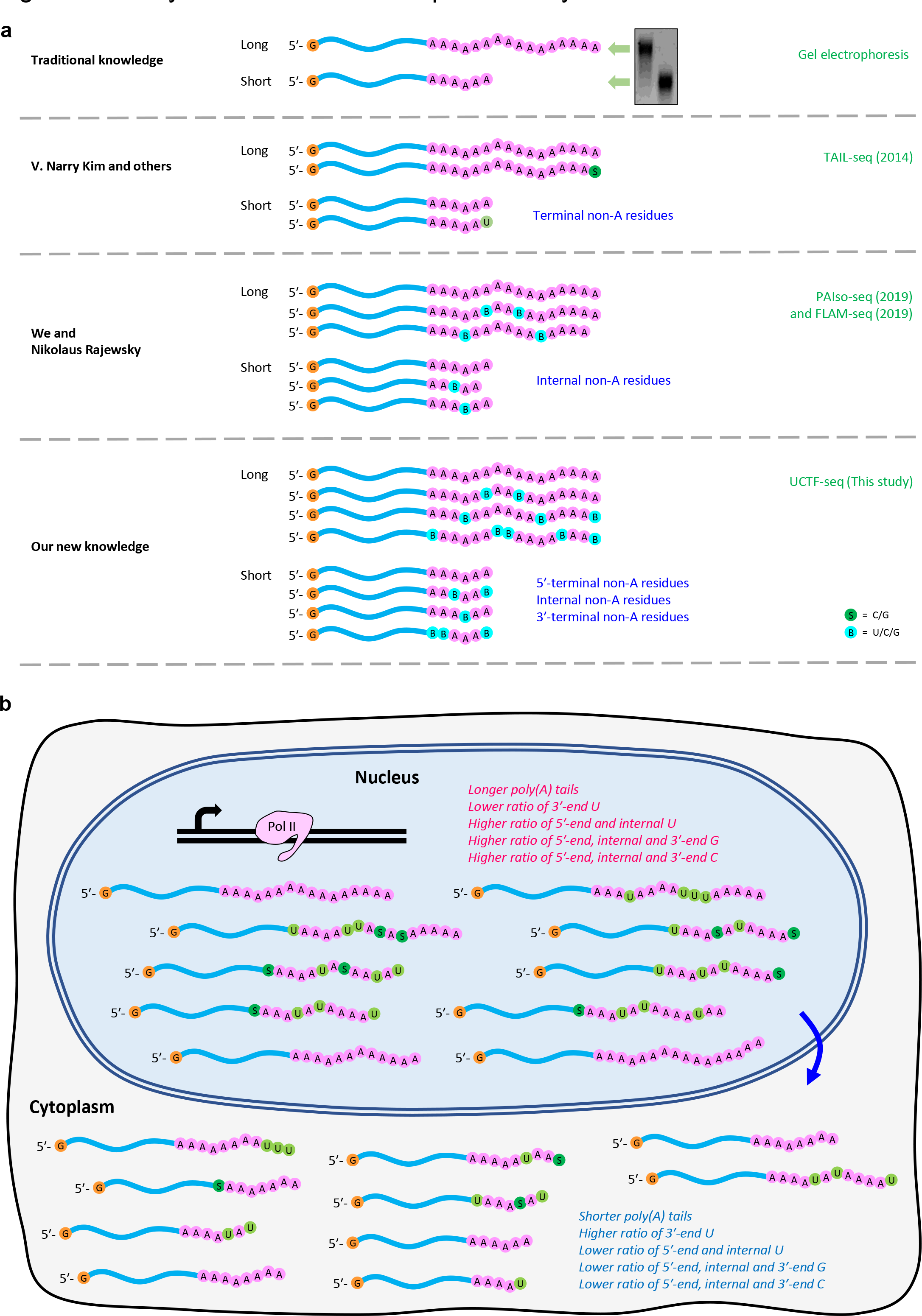
Summary of the mRNA 3′ tail map in a eukaryotic cell. **a,** A historic view of poly(A) tail composition. Traditionally, poly(A) tails were known as pure stretches of A residues of variable length. Then, tail 3′-end non-A residues were discovered by the introduction of TAIL-seq method from the Narry Kim lab. Next, our PAIso-seq together with FLAM-seq from Nikolaus Rajewsky lab revealed wide-spread non-A residues within the body of the poly(A) tails. Now, the UCTF-seq method provides the first comprehensive view of the poly(A) tail base composition, detailing the composition of internal non-A residues as well as those at the 5′- and 3′-ends. **b,** A model for the poly(A) tail composition in the nucleus and cytoplasm. The nuclear RNA has longer poly(A) tails in general. The non-A residues within RNA poly(A) tails have specific features in the nucleus and the cytoplasm.

Hyperadenylation of poly(A)+ RNA has been observed in cells defective in RNA export^31, 32^. However, it remains unknown whether this effect results from a change in subcellular distribution or an indirect effect caused by the nuclear retention of mRNA encoding RNA regulators^34^. Our data reveal that the poly(A) tails of nuclear mRNAs are longer compared to their cytoplasmic counterparts for most genes (Fig. 7b), suggesting that mRNA poly(A) tails undergo extensive processing after nuclear export. In addition to these differences in poly(A) tail length, we also observed biases in poly(A) tails composition in terms of non-A residues, including increased frequencies of 3′-end U residues, and decreased 5′- and 3′-end G residues in cytoplasmic vs nuclear mRNAs (Fig. 7b). The significance of these differences in non-A residue configurations and their link to the changes in poly(A) tail length is an interesting question to be explored in the future.

Our study reveals that the poly(A) tails with internal consecutive U residues may represent a distinct subspecies of mRNAs which is conserved across eukaryotes. Very interestingly, our recent study of mRNA poly(A) tail dynamics during mammalian oocyte-to-embryo transition found that this type of poly(A) tails represents more than 40% of the poly(A)+ transcripts at the 1-cell stage in mouse, pig, rat, and human^35–37^, further suggesting the importance of this new type of poly(A) tails. Therefore, consecutive U residues within the internal body of poly(A) tails define a new type of mRNA poly(A) tails that requires further study. Our ongoing research shows that this new type of tail can pre-mark maternal mRNA for quick degradation at the stage when zygotic genome activation takes place during oocyte-to-embryo transition^35–37^. Canonical polyadenylation polymerases (PAPs) and non-canonical polyadenylation polymerases (ncPAPs) are candidate enzymes responsible for the synthesis of mRNA poly(A) tails^1, 38^. It will be interesting to dissect which ncPAPs and other novel factors contribute to the non-A residues revealed in this study, as well as which factors read the non-A residue information within poly(A) tails.

## Materials and Methods

### Cell culture

3T3 cells were grown in DMEM (Life Technologies) containing 10% fetal bovine serum. Mouse embryonic stem (ES) cells (E14) were grown on 0.1% gelatin-coated plates in DMEM supplemented with 15% fetal bovine serum, penicillin/streptomycin (Life Technologies), nonessential amino acid (Life Technologies), sodium pyruvate (Life Technologies), GlutaMax (Life Technologies), β-mercaptoethanol (Life Technologies), and 1000 U/mL LIF (ESGRO, Millipore). Both types of cells were cultured at 37 °C in a humidified 5% CO2 incubator

### Animals

Mice were purchased from Beijing Vital River Laboratory Animal Technology Co., Ltd and maintained in compliance with the guidelines of the Animal Care and Use Committee of the Institute of Genetics and Development Biology, Chinese Academy of Sciences.

### RNA isolation

Total RNA was extracted with Trizol Reagent (Invitrogen) according to the manufacturer′s instruction. Briefly, samples (total fraction, nuclear and cytoplasmic fractions of 3T3 cells and mouse ES cells (E14), HeLa cell, mouse tissues, rat liver, pig liver, adult zebrafish, adult flies, adult *C. elegans*, *Arabidopsis thaliana*, rice and yeast) were lysed directly in 1 ml of TRIzoland mix thoroughly, then 1 ml of 100% ethanol was added and mixed thoroughly. The mixture was transferred into a Zymo-Spin IC Column (Zymo Research) and centrifuged to capture the RNA on the column. After washing, the RNA was eluted by adding 50 μl RNase-free water directly to the column matrix. The elution step was then repeated. Prepared RNA was stored at -80 ℃ or used immediately.

### Nuclear and cytoplasmic RNA fractionation

Nuclear and cytoplasmic RNA fractionations of 3T3 cells, mouse ES cells (E14) were prepared following a previously published method^27, 28^, and the RNA was further purified using Trizol Reagent (Invitrogen). In brief, cells were harvested and washed with PBS. Then cell pellets were then resuspended in cold lysis buffer (10 mM Tris at pH 8.0, 140 mM NaCl, 1.5 mM MgCl2, 0.5% Igepal, 2 mM vanadyl ribonucleoside complex) and lysed for 5 minutes on ice. One-fifth of the lysate was saved as total RNA fraction. Nuclei were pelleted by centrifuging at 1,000 g for 3 minutes at 4°C. The supernatant was used for cytoplasmic RNA fractionation, and the pellet was used for nuclear RNA fractionation. The pellet was washed twice with lysis buffer followed by a final wash with lysis buffer containing 0.5% deoxycholic acid. After washing, the nuclei were resuspended in a small volume of lysis buffer and nuclear RNA was extracted using Trizol. The fraction containing cytoplasmic RNA was centrifuged at 13,000 rpm for 10 min at 4°C, and the supernatant was transferred carefully to a new tube without disturbing the pellet. The cytoplasmic RNA was further extracted using Trizol.

### UCTF-seq library construction

Total RNA was used for direct 3′ adaptor ligation with T4 RNA Ligase 2, truncated K227Q (NEB) and 3′ adaptor (Supplementary Table 5) at 16°C for 12 hours. The adaptor ligated RNA was cleaned up and concentrated with RNA Clean & Concentrator-5 kit (Zymo Research): the sample was mixed thoroughly with 50 μl RNA binding buffer and 50 μl 100% ethanol; the mixture was transferred into a Zymo-Spin IC Column and centrifuge; after washing, the ligation product was eluted by adding 6- 8 μl nuclease-free water directly to the column matrix and centrifugation. The adaptor ligated RNA was reverse transcribed with template switching method using Superscript II with RT primer and iso-TSO-UMI (Supplementary Table 5). The RT product was used for 1-3 cycles of PCR using PCR primers (Supplementary Table 5). Recombinant Cas9 protein (NEB) and sgRNA prepared by *in vitro* transcription were added to the mixture and incubated at 37℃ for ∼ 4h to remove rRNA. The mixture was cleaned up again with the RNA Clean & Concentrator-5 kit as described above. A second round of PCR amplification was performed using KAPA HiFi HotStart ReadyMix (KAPA Biosystems) with PCR primers. Amplified cDNA products were size selected by Pure PB beads (1x beads for cDNA with sizes above 200 bp and 0.4x beads for cDNA above 2kb). These two parts of the sample were combined at equimolar concentrations for further library construction using SMRTbell Template libraries (SMRTbell Template Prep Kit). The libraries were annealed with the sequencing primer and bound to polymerase, finally the polymerase-bound template was bound to Magbeads and sequenced using PacBio Sequel or Sequel II instruments at Annoroad.

### PAIso-seq library construction

PAIso-seq was performed with total RNA samples according to the previously published method^6^, including templated end extension, reverse transcription, template switching, and PCR amplification. The oligos used for library construction can be found in Supplementary Table 5. The final amplified cDNA products were then size selected by Pure PB beads, and the SMRTbell Template libraries were constructed following UCTF-seq method. The libraries were annealed with the sequencing primer and bound to polymerase, and the polymerase-bound template was bound to Magbeads and sequenced using PacBio Sequel instruments at Annoroad.

### Preparation of DNA and RNA spike-ins

Primers (Supplementary Table 2) were synthesized and annealed to generate double-strand oligos 10A, 28A-GG, 28A-GC, 28A-CC, 29A-C, and 30A. pEASY-T1-Simple vector (TransGen) was treated as pE-0A. The blunt-end oligo 10A was ligated into pE- 0A to generate vector pE-10A. Next, the sticky-end oligo 30A was ligated into pE-10A digested by *Sal*I and *Bsm*BI to generate vector pE-40A. The sticky-end oligos 29A-C, 28A-GC, 28A-GG, 28A-CC, and 30A were ligated into pE-40A digested by *Sal*I and *Bsm*BI to generate vectors pE-70A-C, pE-70A-GC, pE-70A-GG, pE-70A-CC, and pE- 70A. Next, the sticky-end oligo 30A was ligated into pE-70A digested by *Sal*I and *Bsm*BI to generate vector pE-100A. The vector pE-70A-C was digested by *Sal*I and *Bsa*I and the approximately 70 bp oligo was recovered and ligated into pE-70A-GG and pE-70A-CC digested by *Sal*I and *Bsm*BI to generate vectors pE-135A-C/GG and pE- 135A-C/CC, respectively (Extended Data Fig. 1a).

The DNA fragment containing the T7 promoter and mCherry coding region was amplified by PCR with a forward primer containing a *Bam*HI site and reverse primers containing a *Sal*I site and different barcode sequences, to generate the *Bam*HI and *Sal*I flanked T7-mCherry-barcode fragments. Then these fragments were digested by *Bam*HI and *Sal*I, and ligated to pE-0A, pE-10A, pE-40A, pE-70A, pE-70A-GC, pE- 100A, pE-135A-C/GG, or pE-135A-C/CC to generate the template for DNA poly(A) spike-ins 0A, 10A, 40A, 70A, 70A-GC, 100A, 135A-C/CC, and 135A-C/GG. To generate RNA poly(A) spike-ins, *in vitro*transcription using T7 RNA polymerase was performed with PCR product from templates of DNA poly(A) spike-ins 0A, 10A, 40A, 70A, 70A-GC, and 100A (Extended Data Fig. 1a). About 1% of the amount of these spike-ins was mixed to the actual samples for library preparation and sequencing. DNA spike-ins were included in ES-N, ES-C and ES-T runs. RNA spike-ins were included in 3T3-N, 3T3-C and 3T3-T runs.

### sgRNA preparation for rRNA removal

Forward primers containing a T7 promoter, 20 nt variable protospacer sequences targeting mouse rDNA, and a 20 nt sequence pairing with the 5′-end of the sgRNA backbone, and a reverse primer pairing with a 30 nt sequence of the 3′-end of the sgRNA backbone were used to prepare sgRNA DNA template by PCR (Supplementary Table 1). Then PCR was performed with mixed forward primers, reverse primer, a plasmid template containing the sgRNA backbone, and Q5 Hot Start High-Fidelity DNA Polymerase (NEB). After purification, *in vitro* transcription was performed to produce sgRNAs using the HiScribe T7 Quick High Yield RNA Synthesis Kit (NEB). Next, the sgRNAs were cleaned with RNA Clean & Concentrator-5 Kit (Zymo Research) and stored at -80°C.

### UCTF-seq data pre-processing

Circular consensus sequences (CCSs) or reads were generated from subreads using ccs (version 5.0.0), which took multiple subreads of the same SMRTbell molecule and combined them using a statistical model to produce one highly accurate consensus sequence with default parameters. CCS reads provide base-level resolution with > 99.9% single-molecule read accuracy for reads with 10 passes^39^.

To demultiplex and extract the transcript sequence from the CCS reads, we first matched the barcodes in the CCS read with the reverse complement of the CCS reads allowing a maximum of two mismatches or indels (insertions and deletions). CCS reads were oriented and split into multiple transcripts if multiple barcodes were found. Then, to get the precise 3′ end position of the original RNA, we aligned the matched barcode to each transcript using a semi-global function “sg_dx_trace” which did not penalize gaps at both the beginning and the end of query/barcode in parasail package^40^ and trimmed the barcode. Next, we used the following regular pattern “(AAGCAGTGGTATCAACGCAG){e<=2}(AGTAC){s<=1}([ATCG]{8,12})(ATGG G){s<=1}” to match the 5′-adapter and extract the UMIs in each transcript. Finally, the 3′-adapter and the 5′-adapter of each transcript were removed and the remaining extracted sequence was called clean CCS hereafter. Clean CCS reads were used for the downstream analysis.

Clean CCS reads were aligned to the reference genome using minimap2 v.217-r941^41^ with parameters “-ax splice -uf --secondary=no -t 40 -L --MD --cs --junc-bed mm10.junction.bed”. The mm10.junction.bed file was converted from GRCm38 mouse gene annotation with “paftools.js gff2bed” in the minimap2 package. Read counts of each gene and gene assignments of each CCS reads were summarized by featureCounts v2.0.0^42^ with “-L -g gene_id -t exon -s 1 -R CORE -a Mus_musculus.GRCm38.92.gtf” parameters using the read alignments generated by minimap2. Clean CCS reads with the identical mapping position (namely, the start and end position mapped to the reference genome) and the identical UMI (Unique Molecular Identifiers) sequence were determined and only one clean CCS reads was kept. Now the clean CCS reads are ready for downstream analysis.

### PAIso-seq data pre-processing

The demultiplex and extract the transcript sequence from the CCS reads were done following the same procedure as UCTF-seq till the barcode matching steps. Finally, the 3’-adapter and the 5’-adapter of each transcript were removed. Transcripts with length greater than 50nt were retained. Clean CCS reads were used for downstream analysis. Alignment of the clean CCS reads to the reference genome and gene assignment were performed the same way as in the UCTF-seq data pre-processing except that no UMI is involved in PAIso-seq data.

### Poly(A) tail sequence extraction

Clean CCS reads were aligned to the mouse reference genome (mm10) using minimap2 (v.217-r941) with the following parameters “-ax splice -uf --secondary=no -t 40 -L --MD --cs --junc-bed mm10.junction.bed”. Alignments with “SA” (supplementary alignment) tag were ignored. The terminal clipped sequences of the CCS reads in the alignment bam file were used as candidate poly(A) tail sequences. We defined a continuous score based on the transitions between the two adjacent nucleotide residues throughout the 3′-soft clip sequences. To calculate continuous scores, the transition from one residue to the same residue scored 0, and the transition from one residue to a different residue score 1. Number of A, U, C and G residues were also counted in the 3′-soft clip sequences of each alignment. The 3′-soft clip sequences with the frequency of U, C and G greater or equal to 0.1 simultaneously were marked as “HIGH_TCG” tails. The 3′-soft clips which were not marked as “HIGH_TCG” and with continuous score less than or equal to 12 were considered as poly(A) tails.

### Poly(A) tail length measurement

To accurately determine poly(A) tail lengths, we only quantified the poly(A) tail lengths from clean CCS reads with at least ten passes. The poly(A) tail length of a transcript was calculated as the length of the sequence, including U, C or G residues if present. The poly(A) tail length of a gene was presented by the geometric mean of the poly(A) tail length of transcripts with tail length at least 1 nt from the given gene, because poly(A) tail length distribution of a gene is a lognormal-like distribution ^22^.

### Detection of non-A residues in poly(A) tails

Clean CCS reads with at least ten passes were used for calling non-A residues in poly(A) tails. Poly(A) containing U (presented as T in CCSs), C, or G residues were counted as U, C, or G containing transcripts. The percentage of non-A transcripts was the number of poly(A) tails containing at least one G, C or U residues divided by the total number of poly(A) tails with tail length at least 1 nt. The non-A residues at the very 3′-end were defined as 3′-end non-A residues. The non-A residues at the 5′-end while not at the 3′- end were defined as 5′-end non-A residues. The non-A residues within the body of poly(A) tail while not at either 5′-end or 3′-end were defined as internal non-A residues.

### Genome and annotation

The mouse genome sequence used in this study is from the following links. http://ftp.ensembl.org/pub/release-92/fasta/mus_musculus/dna/Mus_musculus.GRCm38.dna_rm.primary_assembly.fa.gz

The mouse genome annotation (including the nuclear encoded mRNAs, lncRNAs and mitochondria encoded mRNAs) used in this study is from the following links. http://ftp.ensembl.org/pub/release-92/gtf/mus_musculus/Mus_musculus.GRCm38.92.gtf.gz

The human genome sequence used in this study is from the following links. http://ftp.ebi.ac.uk/pub/databases/gencode/Gencode_human/release_36/GRCh38.primary_assembly.genome.fa.gz

The human genome annotation (including the nuclear encoded mRNAs, lncRNAs and mitochondria encoded mRNAs) used in this study is from the following links. http://ftp.ebi.ac.uk/pub/databases/gencode/Gencode_human/release_36/gencode.v36.primary_assembly.annotation.gtf.gz

### Data Availability

The ccs data in bam format from PAIso-seq and UCTF-seq experiments generated in this study will be available at Genome Sequence Archive (GSA) hosted by National Genomic Data Center. This study also includes analysis of the following data: Legnini et al. (Gene Expression Omnibus database (GEO): GSE126465), Liu et al. (Sequence Read Archive database (SRA): PRJNA529588), Chang et al. (GEO: GSE51299), and Morgan et al. (ArrayExpress: E-MTAB-5056).

## Acknowledgements

We thank Ying Liu for her technical assistance in cell culturing. We thank Dr. Xiaofeng Cao for the *Arabidopsis* and rice samples, Dr. Qiang Tu for the zebrafish sample, Dr. Zhaohui Wang for the fly sample, Dr. Ye Tian for the *C.elegans* sample, and Dr. Wenfeng Qian for the yeast sample. This work was supported by the Strategic Priority Research Program of the Chinese Academy of Sciences (XDA24020203), the National Key Research and Development Program of China (2018YFA0107001), National Natural Science Foundation of China (31970588, 32170606), Natural Science Foundation of Heilongjiang province (YQ2020C003), the China Postdoctoral Science Foundation (2020M670516, 2020T130687), and the State Key Laboratory of Molecular Developmental Biology.

## Author Contributions

Yusheng Liu, Falong Lu and Jiaqiang Wang conceived the project and designed the study. Yusheng Liu constructed the libraries of the UCTF-seq and PAIso-seq. Yusheng Liu performed all other experiments. Yusheng Liu, Hu Nie, Yiwei Zhang, Falong Lu and Jiaqiang Wang analyzed the sequencing data. Yusheng Liu and Jiaqiang Wang organized all figures. Yusheng Liu, Falong Lu and Jiaqiang Wang supervised the project. Yusheng Liu, Falong Lu and Jiaqiang Wang wrote the manuscript with the input from the other authors.

---

**Extended Data Fig. 1.**
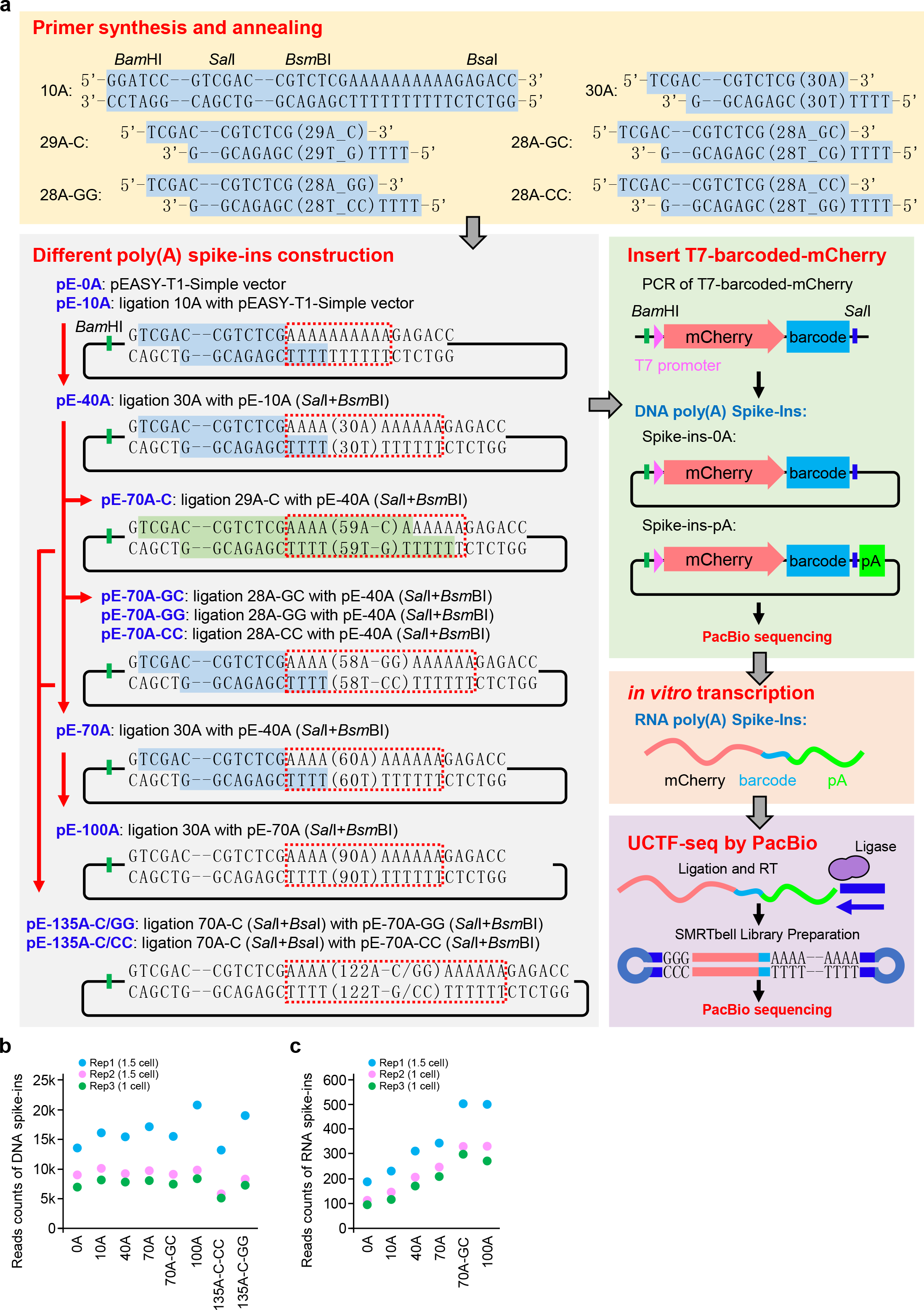
Strategy to generate DNA and RNA poly(A) spike-ins. **a,** Diagram for the preparation of the DNA and RNA poly(A) spike-ins. Strategies for template plasmids construction are shown at the bottom left. The DNA and RNA poly(A) spike-ins used for UCTF-seq are illustrated at the bottom right. Oligos used in the preparation of the DNA and RNA poly(A) spike-ins are shown on the top. **b, c,** Read counts of the DNA spike-ins (**b**) or the RNA spike-ins (**c**) from three experiments. Data points for the three replicates are shown in different colors.

**Extended Data Fig. 2.**
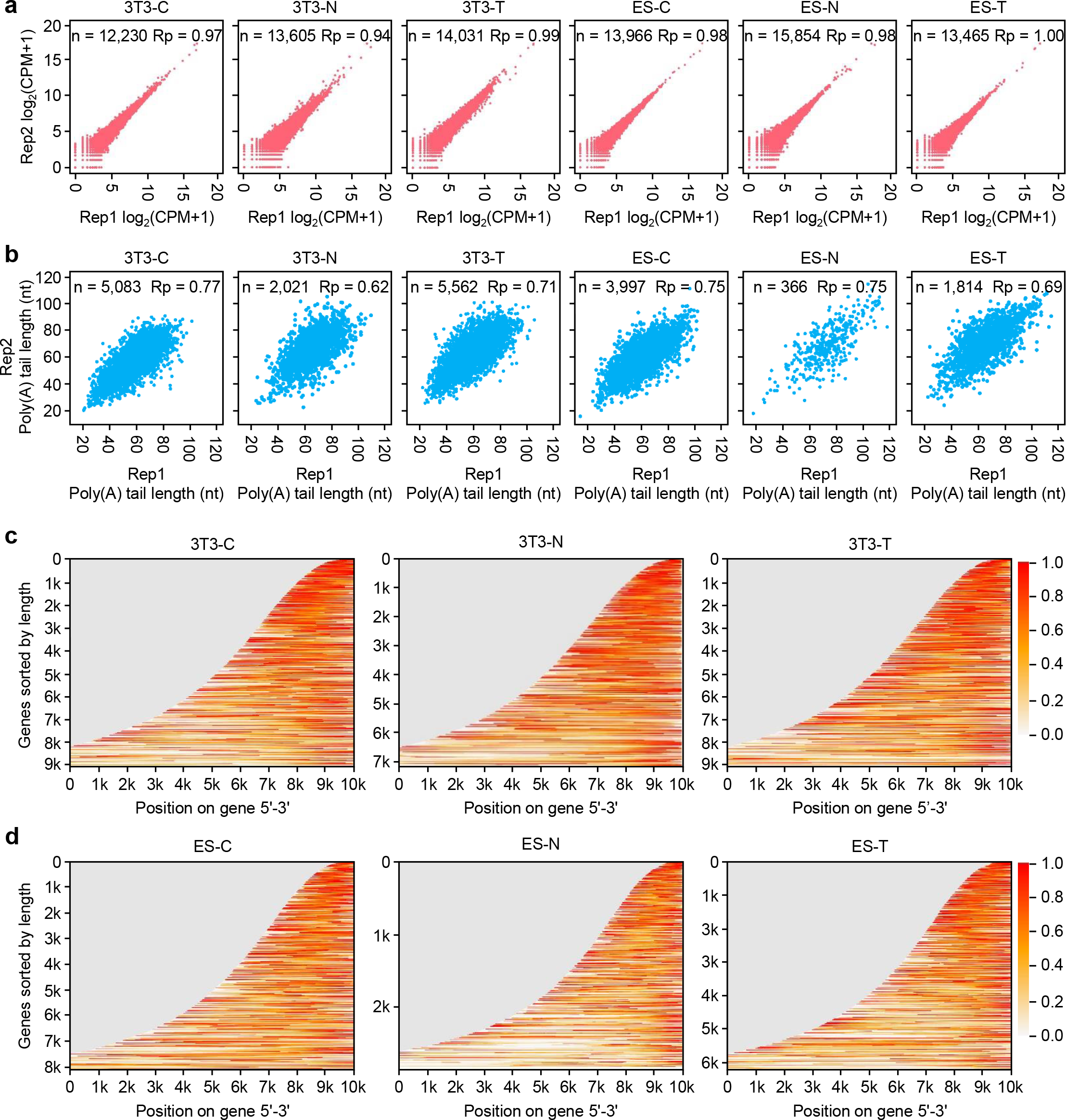
Quality evaluation of the UCTF-seq data. **a,** Correlation of gene expression between two replicates for each sample. The number of genes included in the analysis and Pearson’s correlation coefficient (Rp) of gene expression are indicated for each sample. **b,** Correlation of poly(A) tail length between two replicates for each sample. The number of genes included in the analysis and Pearson’s correlation coefficient (Rp) of gene expression are indicated for each sample. The poly(A) tail length for each gene is the geometric mean length of all the transcripts with poly(A) tail length of at least 1 nt for a given gene. Genes with at least 20 reads in each of the replicates are included in the analysis. **c, d,** Heatmaps showing relative coverage by UCTF-seq across genes in the 3T3 (**c**) or ES (**d**) samples. Genes are ranked by length. The color scale for coverage is included on the right. The X axis represents the length of the genes. The Y axis represents the number of genes included in the analysis.

**Extended Data Fig. 3.**
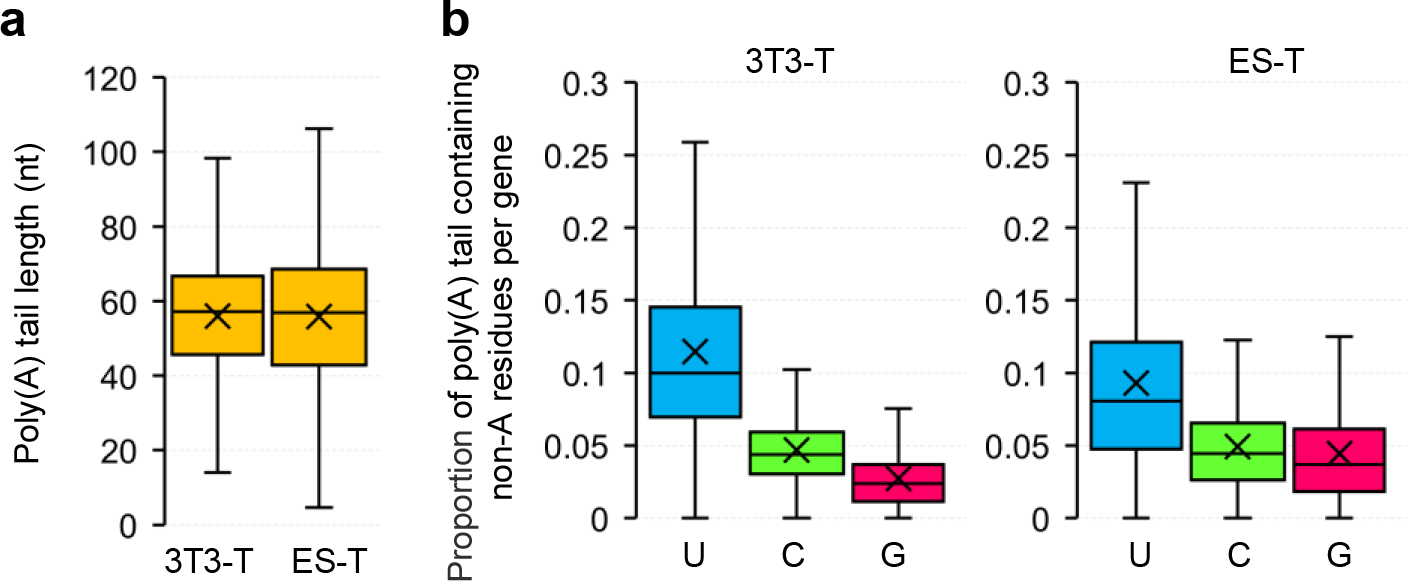
The general characteristics of poly(A) tails in 3T3 and ES samples at the individual gene level. **a,** Box plot for the length of poly(A) tails for each gene in 3T3-T and ES-T samples. The poly(A) tail length for each gene is the geometric mean length of all the transcripts with poly(A) tail length of at least 1 nt for a given gene. **b,** Box plot for the proportion of reads containing U, C or G residues for each gene in the 3T3-T (left) or ES-T (right) sample. The ratio of reads containing U, C or G for each gene is calculated by dividing the number of reads containing U, C or G residues gene by the total number of reads with poly(A) tails of at least 1 nt. For all the box plots, the “×” indicates mean value, the black horizontal bars show the median value, and the top and bottom of the box represent the value of 25^th^ and 75^th^ percentiles, respectively. Transcripts with poly(A) tail of at least 1 nt are included in the analysis.

**Extended Data Fig. 4.**
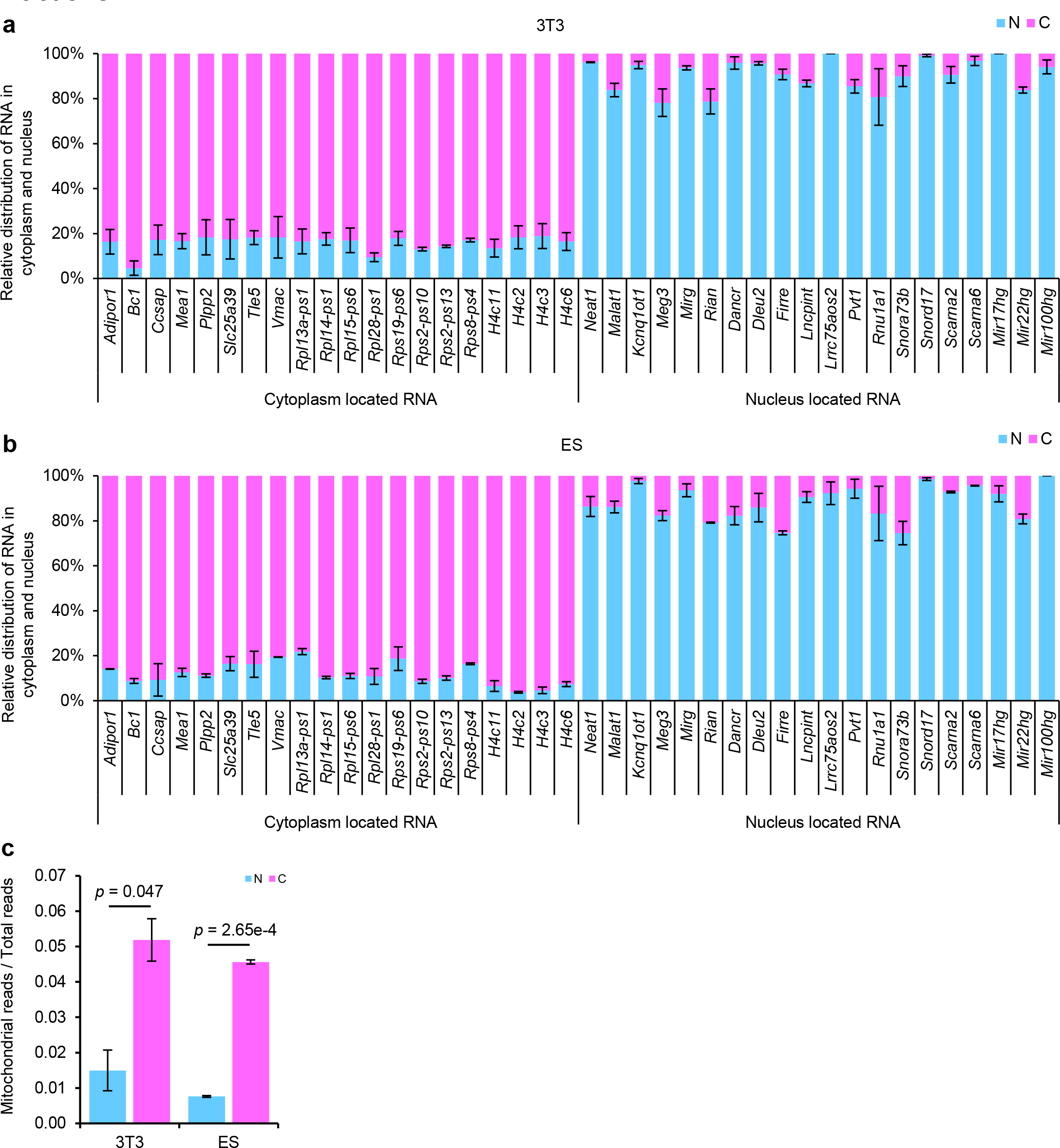
Successful separating the cytoplasmic and the nuclear fractions. **a, b,** Plot of the relative distribution of counts between the 3T3 (**a**) or ES (**b**) nuclear and cytoplasmic fractions of representative genes. The cytoplasmic marker genes are shown on the left, while the nuclear marker genes are shown on the right. **c,** Relative amounts of mitochondria reads sequenced in the nuclear and the cytoplasmic fractions of 3T3 and ES samples. Error bars indicate the SEM from two replicates. All the *p* values are calculated by Student′s *t* test.

**Extended Data Fig. 5.**
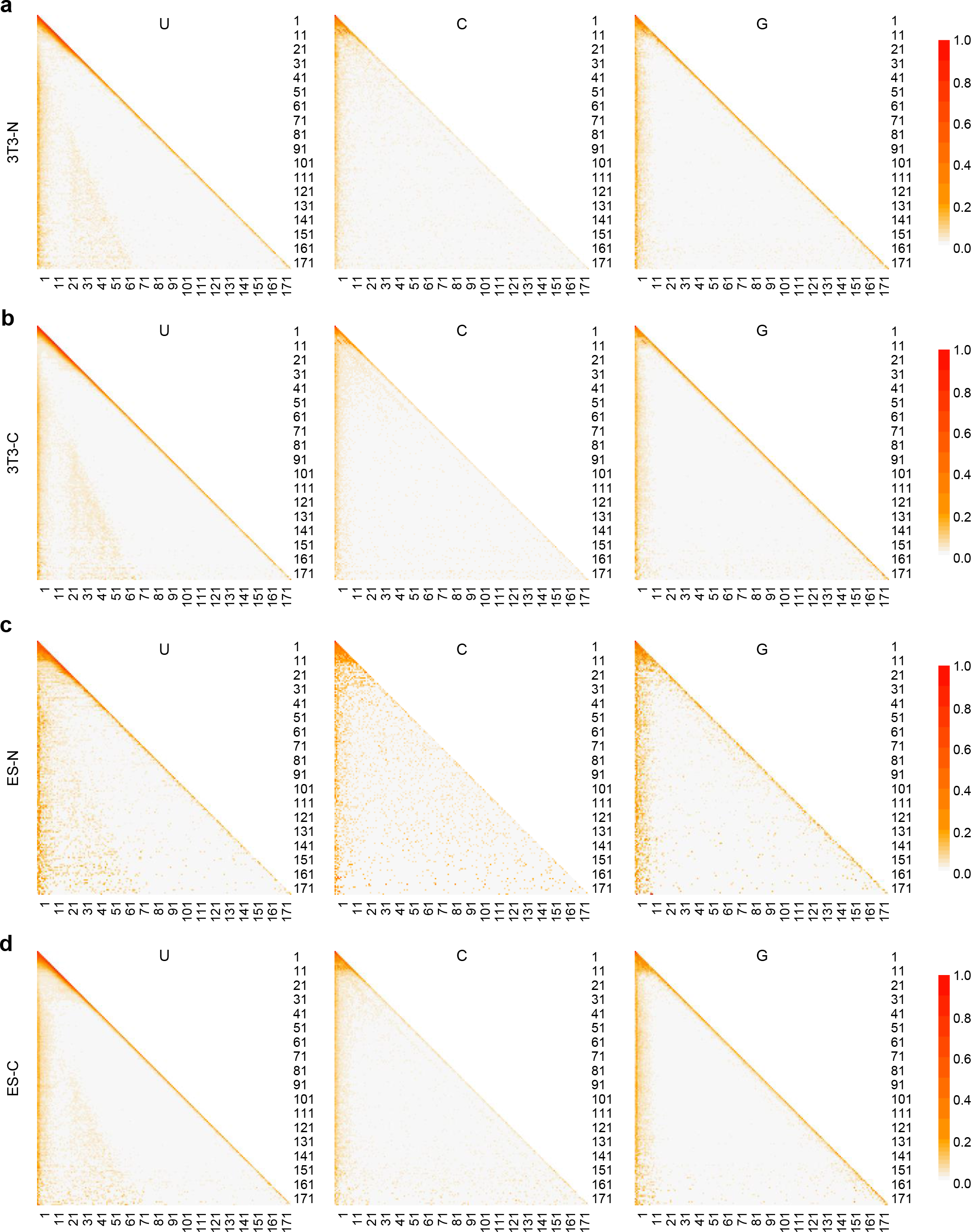
The distribution pattern of U, C or G residues within poly(A) tails in the nuclear and cytoplasmic fractions. Distribution pattern of U, C or G residues within poly(A) tails in the 3T3-N (**a**), 3T3-C (**b**), ES-N (**c**) and ES-C (**d**) fractions. Poly(A) tails with indicated non-A residues of a given length are collapsed to one line. The relative abundance of non-A residues at each position is calculated and visualized by a color scale. Poly(A) tails with length between 1-180 nt are included and ranked in the heatmap from 1-180 (top to bottom).

